# Prediction Dynamically Modulates Gain and Width of Population Frequency Tuning in Human Auditory Cortex

**DOI:** 10.64898/2026.11.16.766995

**Authors:** Jorie van Haren, Ryszard Auksztulewicz, Floris P. de Lange, Sonja A. Kotz, Federico De Martino

## Abstract

During perception, the brain must veridically portray sensory representations while simultaneously highlighting unexpected events. According to two-process theory of predictive perception, this balance is achieved via complementary mechanisms: precision-weighted predictions sharpen sensory tuning around expected features, while momentary surprisal scales response gain, dampening expected inputs and amplifying unexpected violations. However, standard approaches to investigate these processes, such as categorical multivariate decoding, primarily capture shifts in overall signal-to-noise, making it difficult to disentangle changes in population tuning width from modulations in response gain. Here, we combine ultra-high-field functional magnetic resonance imaging (7T fMRI) with population receptive field (pRF) modelling and a continuous Bayesian ideal observer to test how expectations dynamically modulate frequency tuning across the human auditory hierarchy. We first established static pRF preferences and tuning widths independently of predictive inference. We then separately compared this static pRF tuning against two dynamically modulated alternatives. The first model evaluated how tone-by-tone precision-weighted predictions dynamically sharpen and broaden pRF tuning width, while the second model evaluated how gain (by amplification and dampening) was modulated by tone-by-tone information-theoretic surprisal. Across primary, belt, and associative auditory cortices, both a sharpening model driven by precision-weighted predictions and a gain model driven by surprisal explained significantly more variance compared to a static baseline model. Counterfactual models inverting these driving features yielded no improvement, ruling out non-specific model mimicry. Additionally, contrasting local and global tonotopic implementations revealed a feature-selective dissociation: whereas precision-weighted predictions locally sharpened pRFs tuned to expected frequencies, surprisal acted as a global broadcast signal that multiplicatively scaled pRF gain across all frequency channels of the tonotopic map. Together, these findings demonstrate that the auditory system continuously balances selectivity for expected structure and responsiveness to surprising input through local sharpening and global gain modulation.

## 1 Introduction

In the continuous stream of sensory input, the brain faces a fundamental perceptual challenge: it must construct representations that are veridical - reflecting the most likely state of the world - while simultaneously remaining informative by prioritising unexpected events to update internal models. Predictive processing frameworks have traditionally addressed this drive for informativeness by proposing that the brain attenuates predictable information to maintain sensitivity to novelty [1]. However, this cancellation view has been criticised for overlooking the necessity of veridical perception. To reconcile these requirements, a two-process model [2] proposes that probabilistic knowledge initially biases perception toward what is expected, and subsequently upweights events that are particularly surprising (see also [3, 4, 5] for neuromodulatory mechanisms supporting these dynamics). Under this two-process theory, neural tuning properties dynamically adjust as predictive estimates are updated [6]: specifically, tuning-width sharpening (i.e. reduced *σ*) is conceptually linked to the precision of predictions, whereas momentary surprisal drives response gain modulation, dampening expected inputs while amplifying unexpected violations.

Currently, in vivo evidence for such dynamics relies heavily on categorical multivariate approaches - like multivariate pattern analysis (MVPA) decoding - that quantify the representational fidelity of stimuli across different predictive contexts [7, 4, 8]. Within this framework, enhanced decodability (or representational similarity) for expected events is interpreted as evidence for sharpening, whereas higher decodability for unexpected events is interpreted as evidence for gain decrease (dampening) [7, 4]. To clarify what drives these effects, previous work has supplemented multivariate findings with macroscopic univariate observations; however, these attempts have yielded contradictory interpretations. While some studies report that expectation selectively suppresses population-level BOLD activity in a manner consistent with sharpening [9], others find evidence supporting a gain decrease (dampening) account, where suppression is instead greatest in neural populations tuned to the expected stimulus [10]. Because both categorical decoding performance and overall amplitude changes essentially reflect shifts in signal-to-noise ratio, the underlying dynamics at the population tuning level remain mechanistically ambiguous: identical decoding outcomes can arise from entirely distinct neural implementations, such as response gain modulation versus true changes in tuning width [6] (see Figure 1). While recent computational work demonstrates that these conflicting accounts can be resolved by explicitly forward-modelling the underlying population tuning parameters [11, 12, 13, 14], these applications have largely treated these mechanisms as static states across discrete experimental conditions. Conversely, recent electrophysiological work tracking statistical learning demonstrates that tuning-width sharpening and gain modulation are highly dynamic, alternating not only within individual trials but also evolving across distinct learning stages [5]. To fully evaluate the two-process theory, it is therefore necessary to explicitly forward-model both precision-driven sharpening and surprisal-driven gain adjustments and test how each tuning mechanism tracks the evolving predictive context over time.

**Figure 1.**
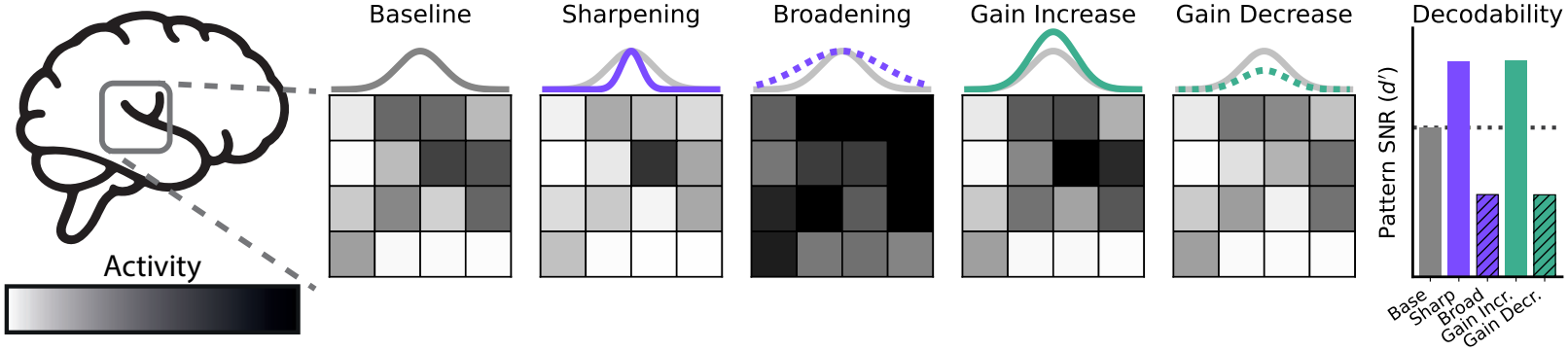
Distinct population tuning mechanisms yield comparable decoding outcomes. Exemplar 1D tuning curves (top) and 4*×* 4 voxel population responses (bottom) for Baseline (grey), Sharpening, Broadening (purple; selectivity modulations), Gain Increase, and Gain Decrease (dampening) (green; amplitude modulations). Right: Multivariate pattern discriminability (*d*^*′*^) between the voxel grids. Sharpening and Gain Increase yield matched increases in *d*^*′*^ above baseline (dashed line), while Broadening and Gain Decrease yield matched decreases, suggesting that macroscopic decoding metrics may obscure differences between changes in tuning width and response gain.

While recent studies have made great strides in tracking predictive dynamics over time [5, 4, 15], resolving how these expectations dynamically reshape sensory selectivity requires moving from categorical condition comparisons to a parametric encoding approach. Sampling stimuli across a continuous acoustic feature space enables the explicit estimation of population receptive fields (pRFs). Combining ultra-high-field (7T) functional MRI with pRF modelling allows us to infer the underlying population tuning mechanisms in a principled manner at the single-voxel level - distinguishing dynamic tuning-width modulations around predicted features from gain modulations driven by unexpected events.

Here we assess how population tuning dynamically adjusts in response to ongoing predictions within an evolving acoustic environment. Utilising stochastic pure-tone sequences with shifting sampling probabilities, we track how predictive processes evolve across frequency space [16] and link these fluctuations to population receptive field (pRF) structure in the auditory cortices using an existing 7T functional magnetic resonance imaging dataset [17]. To investigate the underlying tuning mechanisms, we first establish baseline pRF tuning preference and width in the absence of structured predictions and with minimal short-term adaptation within a localiser session, and validate this static model on the stochastic sequences. We then compare two separate dynamic forward models against this static baseline to assess how hypothesised tuning-width and gain modulations unfold on a tone-by-tone basis, convolving their dynamic predictions with the haemodynamic response and testing their fit to the observed BOLD data. Specifically, we evaluate two mechanistic hypotheses: (i) dynamic tuning-width modulation, where precision-weighted predictions sharpen or broaden pRF tuning width (*σ*_*v*_) around expected inputs, and (ii) dynamic gain modulation, where momentary information-theoretic surprisal (*−*log *P*) scales pRF amplitude via gain increases and decreases. Comparing these models reveals how the auditory system balances selectivity for expected structure with responsiveness to unexpected events.

## 2 Results

Data were previously acquired using ultra-high-field functional magnetic resonance imaging (7T fMRI) over two separate scanning sessions. In the first session, to establish frequency-selective tuning properties independently of predictive inference and stimulus-level adaptation, quasi-random tone sequences were used as functional localisers for population receptive field (pRF) mapping (see Figure 2a). Localiser tone sequences spanned approximately six octaves and were explicitly designed to minimise short-term repetitions and statistical structure. Specifically, successive tones were constrained to differ by at least one octave in frequency. This design minimised temporal correlations between neighbouring frequencies, effectively limiting neural adaptation and improving localised hemodynamic recovery. To model pRF responses, single beta weights were first computed for each voxel, reflecting its activation across the continuum of 240 presented frequencies. Subsequently, a permutation-based model grid search was applied to derive optimal voxel-specific pRF estimates. This permutation approach was used to mitigate the bias toward artificially sharp tuning selectivity that is often observed in regions with lower signal-to-noise ratios [18].

**Figure 2.**
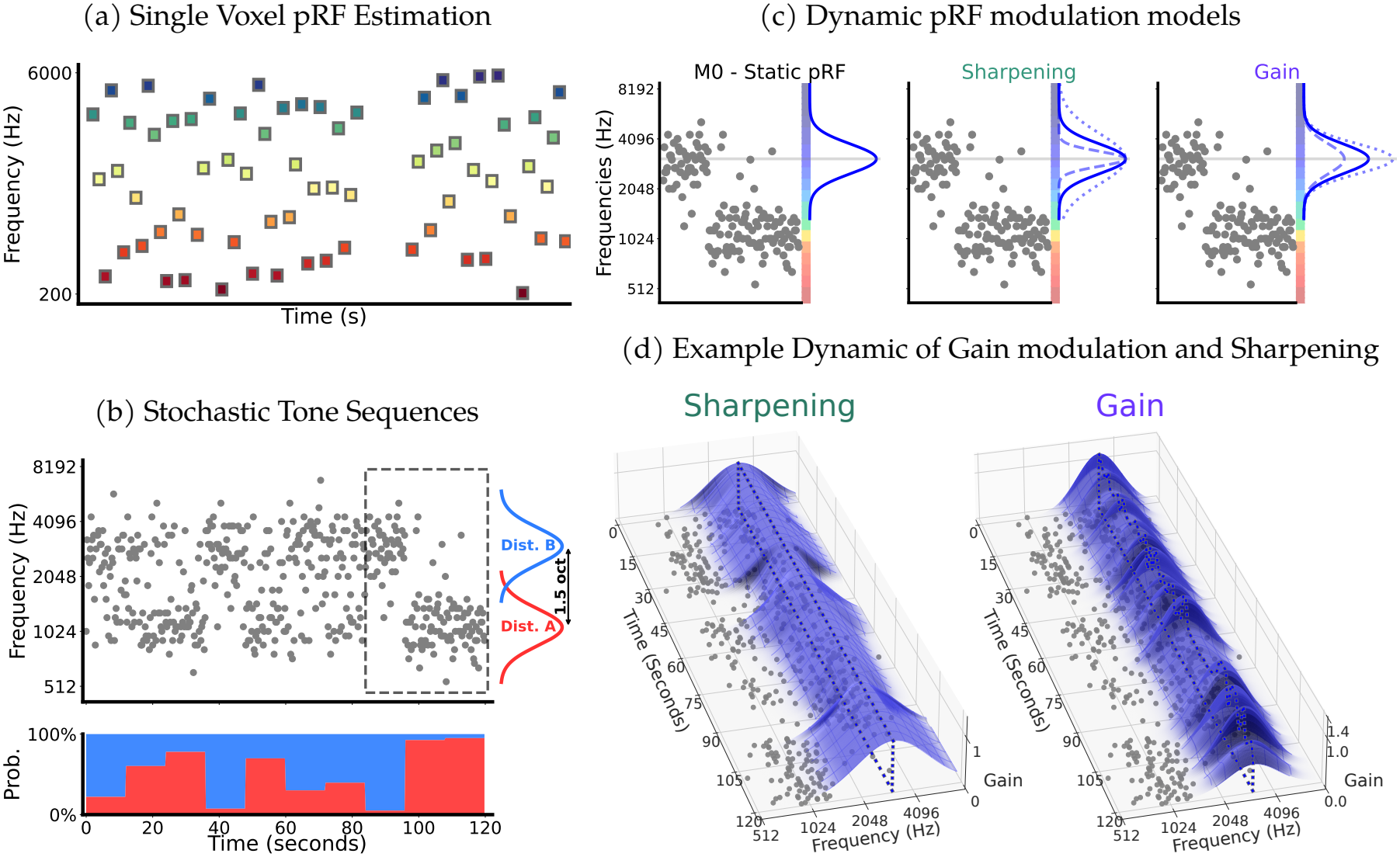
Experimental and Dynamic pRF Modelling Overview. a pRF localiser: Amplitude-modulated pure tones (1.4 s duration, 0.4 s inter-stimulus interval) were presented in quasi-random order, minimising short-term repetition and statistical structure to estimate frequency-selective population receptive fields (pRFs). b Stochastic tone sequences: Pure tones (200 ms duration, 50 ms inter-stimulus interval) were sampled from mixtures of two Gaussian distributions separated by 1.48 octaves. The upper panel shows the presented tone frequencies; the lower panel shows the underlying sampling mixture. c Dynamic adjustments applied to the static (*M*_0_) pRF model estimated from the localiser session. In the sharpening model, the tuning width (*σ*_*v*_) of the pRF is modulated on a tone-by-tone basis, resulting in sharper or broader tuning. In the gain modulation model, the amplitude of the pRF is scaled on a tone-by-tone basis, resulting in amplification or dampening. These modulations continuously update the baseline tuning parameters obtained during the pRF mapping session (example voxel: pRF[3.1 kHz, 0.7 oct]). d Example tone-by-tone trajectories of pRF sharpening (left) and pRF gain modulation (right) for the same tuned voxel over the course of a stochastic sequence, illustrating the distinct predicted activation dynamics under the two separate models.

Functional pRF mapping results, previously detailed in [17], are presented in Figure 3a. Additionally, pRF tuning width estimates are shown in Figure 3b, and individual functional pRF estimates for all ten participants are available in Supplementary Figure S1. Guided by these functional tuning profiles and macroanatomical landmarks (see Figure 3c), we delineated five bilateral regions of interest (ROIs) across the auditory hierarchy: Heschl’s gyrus (HG), Planum Polare (PP), Planum Temporale (PT), and the anterior and posterior superior temporal gyrus (aSTG, pSTG). Within these areas, the vast majority of sound-responsive voxels exhibited robust frequency tuning, surpassing a goodness-of-fit threshold of *r >* 0.2.

**Figure 3.**
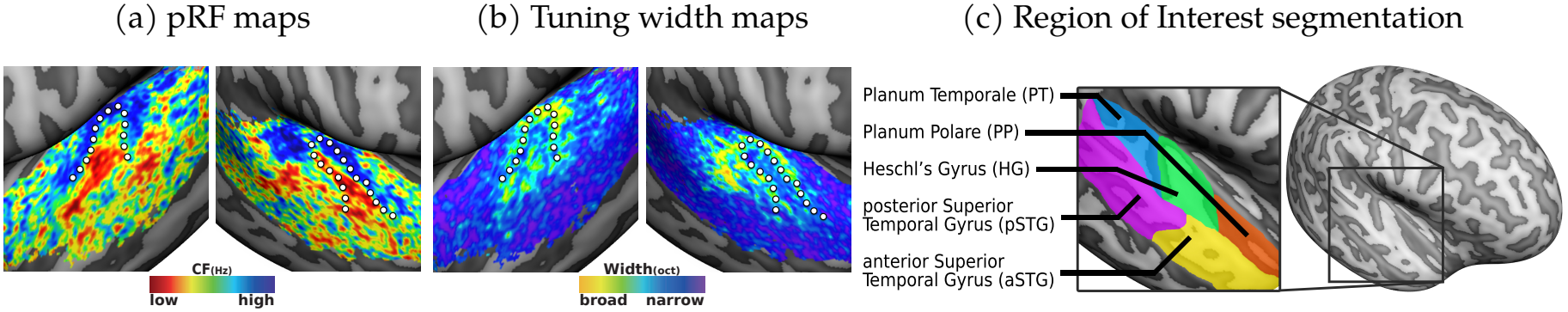
Group-Average Population Receptive Field Maps and Region of Interest (ROI) Segmentation. Group-average pRF maps (*N* = 10, preferred centre frequency; 200 Hz–6 kHz) masked to the auditory cortex and thresholded at *r >* .2. The dotted white line denotes the anatomical border of Heschl’s gyrus. (b) Corresponding tuning width maps (in octaves). (c) Lateral view of the inflated right-hemisphere mid-grey matter surface in a representative subject, with regions of interest delineated according to macroanatomical landmarks [19, 20]. The auditory cortex was subdivided into Heschl’s gyrus (HG), Planum Polare (PP), Planum Temporale (PT), anterior superior temporal gyrus (aSTG), and posterior superior temporal gyrus (pSTG).

Crucially, to ensure these baseline tuning models accurately reflected stable auditory tuning representations, we validated them against independent data from the second scanning session. During this session, participants listened to stochastic auditory sequences of pure tones drawn from Gaussian mixtures with shifting sampling probabilities. Cross-validated regression analyses confirmed that the static baseline pRF model, controlled for low-level effects of repetition suppression, successfully generalised to these stochastic sequences. The established frequency tuning accounted for a significant proportion of the variance in the held-out data across all five regions of interest (all *p*_FDR_ *≤*3.0e *−*3; FDR-corrected; for an overview of all individual statistical tests see Supplementary Table S2). This independent validation confirms that the voxel-wise tuning established in the initial localiser generalises effectively, providing a reliable foundation for testing dynamic predictive modulations.

### 2.1 Evidence for Two-Process Dynamics: Precision Sharpens and Surprisal Scales Auditory Tuning

Having confirmed the stability of the population frequency selectivity across sessions, we next investigated whether BOLD activation patterns are better explained by incorporating dynamic predictive adjustments of population receptive fields. To this end, we separately compared the static pRF baseline model (*M*_0_) against two dynamically modulated alternatives, while controlling for the low-level effects of repetition suppression. In each dynamic model, the static pRF parameters established in the first session served as the structural foundation and were continuously modulated on a tone-by-tone basis over the course of the stochastic sequences. Crucially, rather than adding free parameters to estimate the dynamic fluctuations in tuning width and gain, these tone-by-tone adjustments were used to directly compute the expected activation under the dynamically updating pRFs. By generating a single, voxel-specific expected BOLD time course for each mechanism prior to fitting, this approach ensures that the static baseline and the separate dynamic models maintain identical degrees of freedom.

The first dynamic model evaluated tuning-width modulation, wherein the standard deviation (*σ*_*v*_) of the population receptive field was dynamically adjusted (sharpened and broadened), driven by the localized precision-weighted predictions. The second model evaluated gain modulation, wherein the amplitude of the population receptive field was dynamically scaled, driven by the momentary surprisal elicited by each tone (note that a gain decrease, or dampening is explained by a gain parameter smaller than one). Tone-by-tone predictive quantities driving these models (i.e. precision-weighted predictions and information-theoretic surprisal; *−*log *P*) were derived using a Bayesian ideal observer model (Dynamic Regularity Extraction, D-REX; [16]) that estimates the evolving statistical structure of the tone sequences relying on the preceding acoustic context. We evaluated the evidence present in the data for sharpening and gain modulation separately by testing the increase in out-of-sample variance explained achieved by each dynamic model compared to the static baseline model (*M*_0_).

The dynamic sharpening model, wherein tuning widths were modulated by continuously evolving precision-weighted predictions, significantly increased model accuracy (evaluated in cross- validation) compared to the static baseline (*M*_0_) pRF model throughout the auditory cortices (see Figure 4 and Supplementary Table S3). In the primary auditory cortices, this predictive sharpening resulted in a significant increase in explanatory power over the mean static model (HG: *p*_FDR_ = 2.8e*−*2; proportional improvement 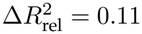). This improvement extended through the auditory belt regions (PP: *p*_FDR_ = 1.5e *−*3, 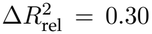; PT: *p*_FDR_ = 6.0e*−* 4, 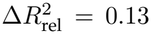) and continued into higher-order associative areas, with additional variance explained in both anterior and posterior portions of the superior temporal gyrus (aSTG: *p*_FDR_ = 1.5e*−* 3, 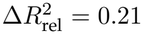; pSTG: *p*_FDR_ = 6.0e *−*4, 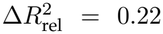). These findings indicate that precision-weighted predictions dynamically adjust (sharpen and broaden) population tuning profiles across the auditory hierarchy. This aligns with computational frameworks proposing that expectations refine sensory representations, extending previous multivariate findings by grounding these effects in explicit models of population receptive field tuning [9, 12, 13].

**Figure 4.**
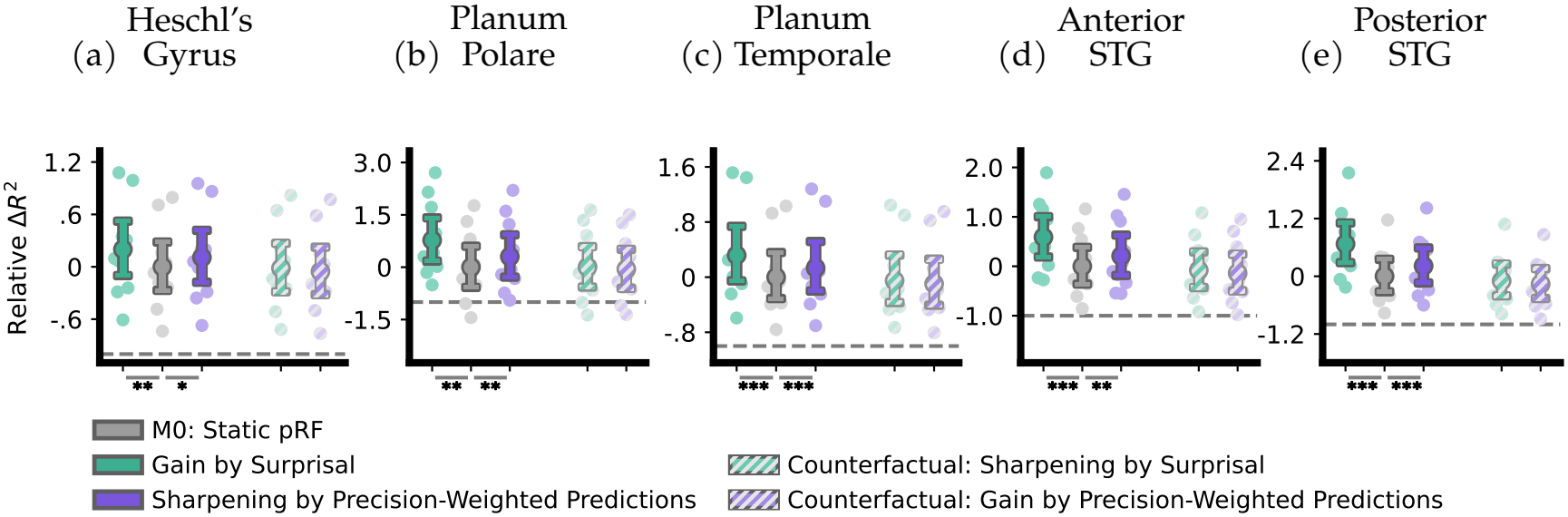
Relative improvement of dynamic modulations over static pRF baseline. Each panel shows the cross-validated relative improvement in variance explained 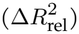 achieved by adding dynamic modulations to the static population receptive field (pRF) model. Results are shown for five auditory regions of interest (ROIs): (a) Heschl’s Gyrus (HG), (b) Planum Polare (PP), (c) Planum Temporale (PT), (d) anterior Superior Temporal Gyrus (aSTG), and (e) posterior STG (pSTG). Primary hypotheses (Gain driven by surprisal and Sharpening driven by precision-weighted predictions) are presented alongside the static pRF model (*M*_0_) to the left of each subplot. Each model’s counterfactual implementations (Sharpening by Surprisal and Gain by Precision-Weighted Predictions) are positioned on the right side of each subplot - distinguished by diagonal stripe patterns. Stars indicate significant relative improvements over the static baseline (FDR-corrected): *p*_FDR_ *<* 0.05 (*), *p*_FDR_ *<* 0.01 (**), *p*_FDR_ *<* 0.001 (***). The horizontal dashed line represents the threshold of zero variance explained (cross-validated *R*^2^ = 0). Data are presented as mean values across participants (*N* = 10) with error bars indicating bootstrapped 95% confidence intervals; all intervals remain strictly above the horizontal dashed line (cross-validated *R*^2^ = 0), confirming out-of-sample predictive power across all models.

Similarly, the dynamic gain model, wherein pRF amplitudes were multiplicatively scaled by the moment-to-moment surprisal elicited by each tone, significantly increased model accuracy (evaluated in cross-validation) compared to the static baseline (*M*_0_) pRF model throughout the auditory cortices (see Figure 4). In primary auditory cortices, this surprisal-driven gain resulted in a significant increase in explanatory power over the mean static model (HG: *p*_FDR_ = 3.0e*−* 3; 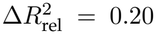). This improvement extended through the surrounding auditory belt regions (PP: *p*_FDR_ = 3.0e *−*3, 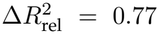; PT: *p*_FDR_ = 3.0e*−* 4, 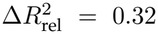) and higher-order associative areas (aSTG: *p*_FDR_ *<* 1.0e*−* 4, 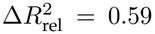; pSTG: *p*_FDR_ *<* 1.0e*−* 4, 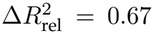). Consequently, surprising auditory events trigger a multiplicative gain increase in pRF responsiveness, whereas predictable events trigger a gain decrease (dampening). This supports the secondary mechanism of the two-process model, wherein unexpected events are upweighted to maintain informativeness [2, 5]. Taken together, these model evaluations demonstrate that precision-driven sharpening and surprisal-driven gain modulation each account for significant dynamic variance over the static baseline throughout the auditory cortical pathways. Finally, evaluating these relative explanatory improvements 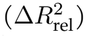 across cortical depth (deep, middle, and superficial compartments) revealed no significant laminar differences for either precision-driven sharpening or surprisal-driven gain across any ROI (all layer-pair comparisons *p*_FDR_*≥* 0.056, with inconclusive Bayes factors ranging from *BF*_01_ = 0.09 to 3.15; for a detailed description of the laminar results, see Supplementary Analysis A.1 and Supplementary Table S1).

### 2.2 Precision Drives Sharpening, Not Gain; Surprisal Drives Gain, Not Sharpening

Having established that both dynamic pRF modulations explain variance beyond the static baseline, a critical remaining question is whether these improvements are specific to their proposed mechanisms. Because our dynamic models integrate higher-level predictive metrics into effective tone-by-tone changes of the pRF (built upon each voxel’s baseline pRF preference), it is possible that incorporating any dynamic predictive factor might artificially capture residual variance and inflate the model fit - a phenomenon known as model mimicry [21, 22, 13]. To explicitly control for model mimicry and ensure our effects are not driven by non-specific correlations, we evaluated counterfactual models in which the driving features were inverted. That is, we tested whether the observed variance could be equally or better explained by precision-weighted predictions driving gain, or by momentary surprisal driving tuning-width sharpening. By directly comparing these counterfactual alternatives against the static baseline, we tested the mechanistic specificity of our primary modulatory effects (for an overview of all counterfactual model performance, see Supplementary Table S3).

We found no evidence that the counterfactual models improved upon the static (*M*_0_) baseline. The precision-driven gain model failed to explain additional variance across any of the auditory regions evaluated (all *p*_uncorrected_ *≥*0.83, see Figure 4, right side of each subplot). To quantify this absence of improvement, we conducted Bayesian Wilcoxon signed-rank tests. These analyses yielded moderate to strong evidence in favour of the null hypothesis (*H*_0_), confirming that the counterfactual models offered no performance gain over the static baseline across Heschl’s gyrus (*BF*_0−_ = 8.39), Planum Polare (*BF*_0−_ = 6.33), Planum Temporale (*BF*_0−_ = 9.67), anterior STG (*BF*_0−_ = 8.46), and posterior STG (*BF*_0−_ = 11.89).

Similarly, the surprisal-driven sharpening model did not perform better than the static (*M*_0_) baseline in any ROI (all *p*_uncorrected_*≥* 0.47, see Figure 4, right side of each subplot). Bayesian analyses again favoured the null hypothesis (*H*_0_) across the auditory hierarchy: HG (*BF*_0−_ = 6.56), PP (*BF*_0−_ = 2.74), PT (*BF*_0−_ = 8.00), aSTG (*BF*_0−_ = 9.08), and pSTG (*BF*_0−_ = 8.10). Together, these counterfactual controls confirm that the two modulatory mechanisms do not mimic one another, demonstrating that within our parametric forward-modelling framework, the improvements in explanatory power are mechanism-specific: precision selectively modulates tuning width, whereas surprisal selectively modulates pRF gain.

### 2.3 Tonotopic Specificity of Predictive Modulations: Global Gain and Local Sharpening

Next, we investigated the feature-selective scope of these predictive modulations across the continuous tonotopic map. Previous computational work in the visual system suggests that predictive sharpening acts locally, selectively sharpening the tuning of populations directly responsive to the expected feature rather than inducing global changes [13]. In contrast, research on neuromodulatory arousal and hierarchical error propagation suggests that surprisal acts as a global broadcast signal, triggering a widespread modulatory response that scales cortical gain across sensory areas [23, 24, 25]. Building on the two-process framework, we explicitly contrasted local and global implementations of our dynamic models to determine whether these respective modulations are restricted to specific frequency-tuned subpopulations or if they apply across the entire tonotopic map (see Figure 5a). Specifically, for precision-driven sharpening, we contrasted a local model - which adjusts tuning widths across the tonotopic map weighted by the full predictive probability distribution (concurrently sharpening expected and broadening unexpected frequencies) - against a global model where the overall precision of the predictions dictates greater, uniform sharpening across all frequencies. Similarly, for surprisal-driven gain, we contrasted a local model - which limits amplitude scaling strictly to the neural populations driven by the current stimulus (boosting only active subpopulations) - against a global model that scales gain uniformly across all frequency channels of the tonotopic map (for a full overview of the results, see Supplementary Table S4).

**Figure 5.**
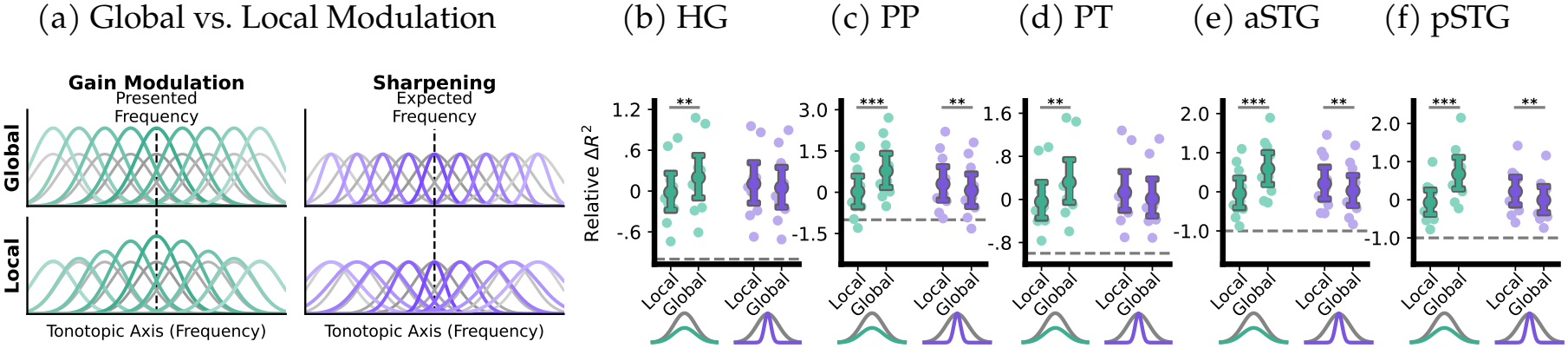
Tonotopic specificity of predictive modulations: Global Gain vs. Local Sharpening. (a) Schematic representations of global and local tuning modulations across the tonotopic axis. Global models uniformly scale gain or sharpen tuning across all frequencies, whereas local models restrict these modulations to the subpopulations tuned to the presented (gain) or expected (sharpening) frequencies. (b–f) Cross-validated relative improvement in variance explained 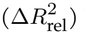 between local and global implementations of the dynamic predictive models (above static baseline *M*_0_). Results are shown across five auditory regions of interest (ROIs): Heschl’s Gyrus (HG), (c) Planum Polare (PP), (d) Planum Temporale (PT), (e) anterior Superior Temporal Gyrus (aSTG), and (f) posterior STG (pSTG). Within each panel, the left cluster (green) compares local versus global implementations for surprisal-driven gain modulation, and the right cluster (purple) compares local versus global implementations for precision-driven sharpening. Schematic Gaussians below the axes illustrate the respective dynamic mechanisms. Significance bars indicate FDR-corrected comparisons between the local and global implementations: *p*_FDR_ *<* 0.05 (*), *p*_FDR_ *<* 0.01 (**), *p*_FDR_ *<* 0.001 (***). The horizontal dashed line 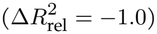 represents the threshold of zero absolute variance explained (raw cross-validated *R*^2^ = 0). Data are presented as mean values across participants (*N* = 10) with error bars indicating bootstrapped 95% confidence intervals; all intervals remain strictly above the horizontal dashed line, confirming out-of-sample predictive power across all models.

For surprisal-driven gain modulation, we found that a global implementation consistently outperformed the local model (see Figure 5b–f). Global gain modulation provided a significantly better fit in the primary auditory cortex (HG: *p*_FDR_ = 1.5e*−* 3; 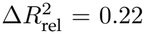), the auditory belt (PP: *p*_FDR_ = 1.0e *−*4, 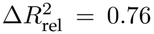; PT: *p*_FDR_ = 1.5e*−* 3, 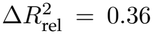), and associative regions (aSTG: *p*_FDR_ = 3.0e*−* 4, 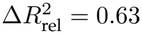; pSTG: *p*_FDR_ *<* 1.0e*−* 4, 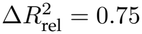). This indicates that the momentary surprisal elicited by a tone uniformly scales population responsiveness across the broader tonotopic landscape. This widespread amplification is consistent with the hypothesis that surprisal operates as a global broadcast signal, potentially mediated by the widespread projections of neuromodulatory arousal systems [23].

Conversely, precision-driven sharpening was better characterised by a local implementation, though the robustness of this effect varied across the auditory hierarchy (see Figure 5b–f). The local sharpening model significantly outperformed its global counterpart in parts of the auditory belt (PP: *p*_FDR_ = 5.9e*−* 3; 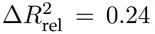) and associative regions (aSTG: *p*_FDR_ = 3.6e*−* 3, 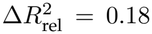; pSTG: *p*_FDR_ = 3.6e*−* 3, 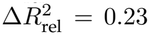). Throughout the remaining areas of the auditory cortex, the local model consistently performed better numerically but did not reach FDR-corrected significance, specifically in the primary auditory cortex (HG: *p*_FDR_ = 3.8e*−* 1; 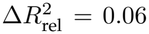) and the remaining auditory belt (PT: *p*_FDR_ = 6.2e*−* 2; 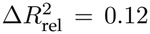). Furthermore, Bayes factors indicated little to no evidence supporting the null hypothesis in these areas (*BF*_01_ = 1.20 and *BF*_01_ = 0.10, respectively; notably, both local and global models still significantly outperformed the static baseline everywhere).

Together, these results suggest that precision-driven tuning-width modulation operates as a context-dependent filter, albeit with regional differences in statistical robustness. By selectively sharpening the tuning of neural populations centred on the expected frequency, the auditory system locally enhances selectivity for predicted features while simultaneously broadening tuning for uninformative contexts [13, 14].

## 3 Discussion

In this study, we investigated how tone-by-tone auditory expectations dynamically shape population receptive field (pRF) tuning across the human auditory hierarchy. By combining ultra-high-field (7T) fMRI with computational modelling of inferential estimates, we evaluated core assumptions of the two-process framework of predictive perception [2]. Specifically, we used normative estimates derived from a Bayesian ideal observer to continuously modulate baseline tuning parameters in separate forward models, testing whether precision-weighted predictions drive local tuning-width modulation and momentary surprisal drives gain modulation. Our findings indicate that predictive computations modulate neural responses through two dissociable mechanisms: precision-weighted predictions locally sharpen the tuning of neuronal populations associated with expected frequencies, whereas momentary surprisal acts as a global multiplicative factor that scales pRF gain uniformly across all frequency channels of the tonotopic map.

These results offer univariate, population-level support for the two-process theory, expanding upon the mechanistic ambiguities often present in widely used multivariate decoding approaches [6, 12, 5]. By explicitly forward-modeling the underlying ongoing tuning modulation, we established that precision-weighted predictions and surprisal subserve distinct modulatory roles. The selective sharpening of pRFs driven by precision-weighted predictions operates as a context-dependent filter. By sharpening the tuning of populations explicitly associated with expected frequencies, the auditory system enhances selectivity for predicted features while broadening tuning for uninformative contexts [13, 14]. Conversely, the global nature of the gain model demonstrates that momentary surprisal does not act in the same context-dependent manner. Rather, it acts as a widespread, context-uninformative broadcast signal alerting the system that prevailing expectations have been violated [26]. In line with two-process accounts [2], this global mismatch interrupts the brain’s initial bias toward expected structure, facilitating the evaluation of the exact nature of the deviation and potentially prompting a shift into an active ‘exploration’ state [27, 28].

Furthermore, by linking population receptive field modulations to a continuously updating Bayesian observer model, our results highlight the dynamic nature of population receptive fields. Previous studies investigating tuning curve modulation - whether driven by expectations or attention - have primarily compared population tuning parameters across discrete experimental blocks [29, 11, 12, 13, 30, 14]. Extending this work to a continuously evolving stochastic soundscape, we demonstrate that tone-by-tone fluctuations in sharpening driven by precision-weighted predictions, and gain driven by surprisal, each capture significant BOLD variance beyond static frequency selectivity. Guided by the ongoing acoustic context, this establishes that the perceptual advantages of predictive processing extend directly to the sensory selectivity of neural populations, resulting in a continuous structural reshaping of how the cortex represents sensory feature input.

While our dynamic modelling approach allowed us to test specific, hypothesis-driven predictions regarding tuning width and gain, these inferences remain bounded by the mesoscopic resolution of high-field fMRI. Spatially, because our approach relied on the tonotopic organisation of the cortex to deduce univariate population codes, the observed dynamics fundamentally reflected the aggregate behaviour of tens of thousands of neurons. Consequently, it remains possible that the macroscopic sharpening observed here emerges from localised, feature-specific gain modulation and dampening at the level of individual neurons [6]. Furthermore, temporally, the sluggishness of the haemodynamic response limits our ability to resolve the rapid dynamics suggested within the two-process framework. Both theoretical accounts and recent electrophysiological evidence demonstrate that the initial bias toward expected structure and the subsequent upweighting of surprising inputs are separable in time, unfolding as a rapid two-phase process during individual trials [2, 5]. Because fMRI integrates signal over seconds, our measurements inherently aggregate these temporally distinct processing stages, capturing ongoing sequence-level tuning reshaping rather than trial-level mechanisms. To fully capture how these rapidly alternating mechanisms evolve while preserving the advantages of univariate analysis, future research could translate this explicit pRF modelling framework to invasive recordings (e.g., intracranial electroencephalography), or to neuroimaging modalities that combine high temporal precision with distinguishable population tuning profiles (e.g., using parametric tuning estimation [12]). Lastly, beyond these spatial and temporal considerations, it remains to be tested whether these mechanisms generalise from pure tones to the more complex acoustic features representations present in higher-order auditory regions.

Together, our findings demonstrate that the human auditory cortices continuously balance veridical and informative perception by deploying two complementary tuning strategies. Precision- weighted predictions locally sharpen sensory representations, while unexpected events globally scale cortical gain to facilitate exploration of the acoustic environment. Operating across distinct local and global scopes, these spatially concurrent but computationally distinct mechanisms allow the auditory system to maintain selectivity for structured acoustic environments without sacrificing sensitivity to novel information.

## 4 Methods

The computational and analytical framework presented here builds upon ultra-high field fMRI data (at 7T) previously acquired and detailed in [17]. The current study introduces a novel dynamic population receptive field (pRF) modelling approach. By leveraging previously established tonotopic maps, we directly estimate and test temporal fluctuations in population receptive field (pRF) parameters induced by predictive inference.

### 4.1 Participants and Ethical Approval

Ten participants (five female; *M* = 22.6 years, age range: 18–29 years) with normal hearing and no history of neurological disorders took part in the fMRI experiment. The study was approved by the Ethics Review Committee of the Psychology and Neuroscience faculty (ERCPN; Approval Code: OZL-232-01-01-2021) at Maastricht University and was conducted in accordance with the Declaration of Helsinki. Written informed consent was obtained prior to the experiment, and participants received monetary compensation.

### 4.2 MRI Acquisition and Stimuli Delivery

MRI data were acquired on a 7-Tesla Siemens MAGNETOM scanner using a 32-channel head coil (Nova Medical) at Scannexus (Maastricht, The Netherlands). All auditory stimuli were generated in MATLAB using the Psychophysics Toolbox [31, 32] and delivered via MRI-compatible in-ear headphones (Sensimetric Inc., s15 model) at a 48,000 Hz sampling rate. Prior to scanning, a loudness calibration procedure (using 9 tones spaced between 200 and 6,000 Hz) was used to construct individual equal-loudness contours, ensuring uniform perceived intensity across the frequency spectrum [33].

During Session 1, high-resolution T1-weighted anatomical data were collected (0.7 mm isotropic, MP2RAGE), alongside functional MRI data acquired with a 2D gradient-echo (GE) simultaneous multi-slice (SMS) EPI sequence (TR = 1800 ms, TE = 26 ms, 0.8 mm isotropic, MB = 2, GRAPPA = 3). Session 2 utilised identical functional acquisition parameters, with slice positioning automatically aligned to the first session. Collectively, these sessions resulted in approximately 4 hours of functional scanning time per participant.

### 4.3 Population Receptive Field Mapping (Session 1)

To establish a baseline model of auditory frequency selectivity independent of sequence structure, Session 1 mapped population receptive fields (pRFs) across the auditory cortex using quasi-random tone sequences.

#### 4.3.1 Quasi-Random Tone Sequences

We measured fMRI responses to sequences of 240 pure tones spanning approximately six octaves. Each tone was presented once per run (duration: 1.4 s, ITI: 400 ms). To allow for the unbiased estimation of frequency selectivity, the presentation order was quasi-randomised with the requirement that successive tones differed by at least one octave. This constraint minimised temporal correlations between neighbouring frequencies, effectively limiting neural adaptation and improving localised hemodynamic recovery. Session 1 included six pRF mapping runs (approximately 8 min 20 s each), though data corruption limited one participant to five usable runs. To stabilise baseline estimations and promote recovery, 14.4-second silent intervals were inserted after every 80 tones, and brief 1-TR silent gaps were randomly distributed across 5% of trials.

#### 4.3.2 pRF Estimation and the Static Baseline Model (*M*_**0**_)

pRFs were estimated using a two-stage procedure. First, voxel-wise single beta weights were computed for each of the 240 frequencies relying on a canonical double-gamma hemodynamic response function (HRF). Second, a permutation-based grid search (1,000 permutations) was applied to derive optimal voxel-specific pRF estimates (*µ*_*v*_, *σ*_*v*_) while mitigating the bias toward artificially sharp tuning often observed in low-SNR regions [18, 17]. The resulting predicted response *R*_*v*_ to a tone frequency *x*_*t*_ was defined as a Gaussian response profile in log_2_-frequency space:

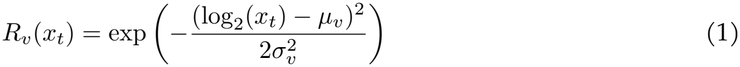

Crucially, this static formulation constitutes our baseline model (*M*_0_). For any given tone presented in Session 2, the expected activation under *M*_0_ is the height of the static 1D Gaussian at that tone’s frequency. This serves as the foundation against which all dynamic temporal modulations are generated and compared.

### 4.4 Presentation of Stochastic Sequences (Session 2)

To manipulate short- and longer-term predictive inference, Session 2 employed stochastic tone sequences sampled from Gaussian mixtures (bandwidth: 1/3 octave FWHM). The relative sampling probability from these distributions shifted periodically (ranging from strongly biased 95%/5% mixtures to more balanced 55%/45% mixtures). Consequently, the predictive inference landscape associated with this context evolved continuously over time. This dynamic design was explicitly chosen to dissociate sequence-level expectation from repetition suppression by continuously fluctuating local statistical regularities.

### 4.5 Functional Data Analysis

#### 4.5.1 Preprocessing and Region of Interest Selection

Functional data preprocessing included slice timing correction, 3D rigid-body motion correction, temporal high-pass filtering (7 cycles/run), Topup geometric distortion correction, and ANTs non-linear alignment between scanning sessions.

Based on macroanatomical landmarks and reliable frequency selectivity established during pRF mapping, we delineated five bilateral regions of interest (ROIs): Heschl’s gyrus (HG), Planum Polare (PP), Planum temporale (PT), anterior superior temporal gyrus (aSTG), and posterior superior temporal gyrus (pSTG).

For supplementary analyses evaluating model performance across cortical depth (deep, middle, and superficial; see Supplementary Analysis A.1), the cortical grey matter was partitioned using an equi-volume approach [34], assigning each functional voxel (0.8 mm isotropic) to a single depth compartment based strictly on majority volume overlap with the high-resolution anatomical segmentation (0.4 mm isotropic).

### 4.6 Computational Modelling of Predictive Features

#### 4.6.1 Predictive Inference (D-REX)

To simulate the temporal evolution of auditory expectations, we employed the Dynamic Regularity Extraction (D-REX) model [16]. This Bayesian observer continuously predicts the distribution of the next stimulus *x*_*t*+1_ based on previous observations *x*_1:*t*_ by estimating local statistics *θ* (the sample mean and variance of the input). To resemble perceptual constraints, model memory (*m*) was restricted to remain within the boundaries of a single block (the 2-minute stimulus period between silent pauses).

To represent multiple components within the sound sequences, the model utilises a Gaussian Mixture Model (GMM). Given our stimulus design, we constrained the maximum number of GMM components to two. To manage the formation of new components, a sparsity hyperparameter (*β* = 0.2) was applied to the mixture weights, encouraging the model to concentrate probability mass into fewer components unless a highly diverging stimulus occurs.

D-REX accounts for potential environmental volatility by generating hypotheses across varying run lengths (*r*_*t*_), where *r*_*t*_ represents the number of observations since the last change point. Each hypothesis is assigned a belief equivalent to its posterior probability, *P* (*r*_*t*_ | *x*_1:*t*_). These context-specific predictions are then weighted by their respective beliefs and integrated to predict future observations:

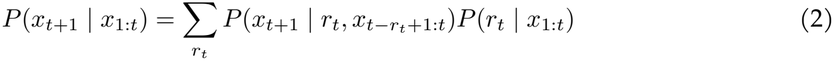

From this integrated predictive distribution, we extracted two key computational metrics for every tone presentation:

Surprisal (*S*_*t*+1_), representing the discrepancy between the predicted probability and the actual observation, where lower probability events yield higher surprisal:

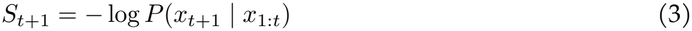

Precision and Precision-Weighted Predictions, where the model computes global sequence precision as the inverse of the variance across the full Gaussian mixture predictive distribution, quantifying overall contextual confidence:

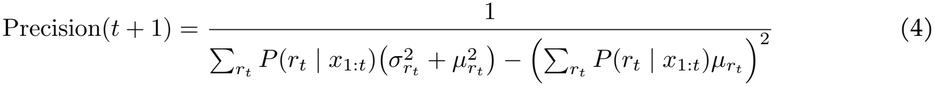

Crucially, because D-REX generates a continuous probability distribution across the entire stimulus space, we also extracted the localised predictive probability mass across the frequency spectrum. This serves as a *precision-weighted prior*, explicitly detailing the model’s expectation for specific frequency channels at any given moment, enabling localised tuning adjustments.

#### 4.6.2 Repetition Suppression Control

To account for the persistent, stimulus-specific suppression induced by brief sensory exposure, all model activations were multiplicatively weighted by a double-exponential long-trace repetition suppression model prior to analysis. Based on previously established and verified parameters [35, 17], this integrated both fast and slow adaptation components reflecting the sequence’s specific stimulus history, ensuring that predictive models were evaluated independently of low-level adaptation mechanisms.

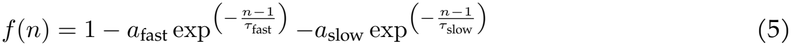

where *a*_fast_ = 13.99% and *a*_slow_ = 3.45% determine the amplitudes, and *τ*_fast_ = 0.85 tones and *τ*_slow_ = 6.82 tones are the respective time constants for the fast and slow suppression components [35].

### 4.7 Dynamic pRF Modulation

Our primary objective was to investigate the mesoscopic neural implementation of predictive processing by explicitly parameterising two distinct adjustments to our static *M*_0_ baseline.

#### 4.7.1 Dynamic Gain Modulation

To test whether predictive context scales the overall amplitude of neural responses without altering selectivity, we introduced a dynamic gain factor driven by Surprisal:

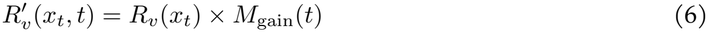

To explicitly capture both predictive gain decreases (dampening) and expectation-driven gain increased, our generative models computationally constrained this scaling multiplier to a bounded range between 0.5 and 1.5 using a logistic transition. Within this continuous axis, a highly predictable, low-surprisal context yields an *M*_gain_(*t*) *<* 1 (gain decrease), while a highly surprising event yields an *M*_gain_(*t*) *>* 1 (gain increase). Because the resulting time courses are fitted to *z*-scored fMRI data using Ordinary Least Squares (OLS) regression with an intercept, the critical feature is the dynamic - proportional fluctuations - of response amplitudes tracking surprisal.

#### 4.7.2 Dynamic Tuning-Width Modulation (Sharpening)

To test whether the brain optimises encoding efficiency by sharpening or broadening neural tuning around expected inputs, we dynamically modulated the voxel’s spatial standard deviation (*σ*_*v*_). Under the primary hypothesis, this modulation in sharpening and broadening (*σ*_*v,t*_) is driven locally by precision-weighted predictions:

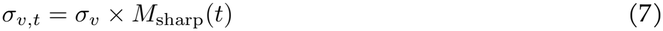

where *M*_sharp_(*t*) is a multiplier derived from the predictive probability at that specific frequency. High predictive probability yields *M*_sharp_(*t*) *<* 1 (sharpening up to a limit of 0.5*σ*), whereas low probability yields *M*_sharp_(*t*) *≥* 1 (broadening up to 1.5*σ*). This time-varying *σ*_*v,t*_ replaces the static *σ*_*v*_ in Equation 1.

#### 4.7.3 Model Comparisons and Spatial Scope

First, each dynamic model was compared separately against the static baseline pRF model (*M*_0_) to determine whether it explained additional variance beyond static frequency selectivity.

Next, to evaluate the evidence for the specific proposed mechanisms and control for model mimicry, we evaluated counterfactual models where the driving features were inverted (e.g., models where precision-weighted predictions drove Gain Modulation, and Surprisal drove Tuning-Width Sharpening).

Finally, because predictive metrics can represent general contextual states or feature-specific estimates, we investigated the spatial scope of each model. We compared models having a *Global Scope*, where the modulation factor is applied uniformly across the entire tuning grid (e.g., a highly surprising event globally upweights gain across all frequency channels regardless of preferred frequency), against models having a *Local Scope*, where the modulation factor is functionally restricted to specific frequency channels (e.g., tuning is sharpened strictly for the neuronal populations tuned to the currently predicted frequencies).

### 4.8 fMRI Regression and Statistical Analysis

All tone-by-tone dynamic predictions were generated at a 50 ms resolution, adjusted for repetition suppression, convolved with a canonical double-gamma HRF, and resampled to the fMRI repetition time (TR = 1.8 s). Analyses were restricted to sound-responsive voxels, defined as those demonstrating a significant in-sample fit (*p <* 0.05) for a general sound ‘on’ vs ‘off’ period, as determined by an F-test evaluating the overall variance explained by the model’s predictions.

Voxel-wise model performance was evaluated using a leave-one-out cross-validated *R*^2^ regression approach. The structural simplicity of the predictors allows models to be composed of identical, single-vector stick predictors representing expected activations over time under the different tuning Gaussians. That is, by comparing the cross-validated variance explained by the static baseline model against its dynamically modulated counterparts, we directly isolate the explanatory power of specific theoretical mechanisms.

Statistical significance was assessed at the region-of-interest (ROI) and cortical-layer level across participants (*N* = 10) using non-parametric bootstrap *t*-tests (10^5^ iterations). To control for multiple comparisons, p-values were corrected using the False Discovery Rate (FDR) via the Benjamini–Hochberg procedure across all ROIs and depth compartments.

To quantify evidence for the absence of effects in counterfactual models that control for model mimicry, we supplement our non-parametric analyses with non-parametric Bayes Factors. Specifically, we computed Bayesian Wilcoxon signed-rank tests using a default Cauchy prior (scale 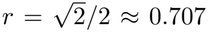) on the latent (standardised) effect size, estimated via data augmentation with 10,000 samples [36]. For these controls, we report the directional *BF*_0*−*_ to indicate the relative evidence for the null hypothesis (i.e., no improvement in explained variance compared to the static baseline) over the alternative.

## Acknowledgements

We thank Agustin Lage-Castellanos for his implementation of the permutation testing for population receptive field modelling.

## Data Availability

The anatomical and functional MRI data used in this study have been fully anonymised, defaced to ensure participant privacy, and deposited in the OpenNeuro database under accession code ds006928 [37] (https://openneuro.org/datasets/ds006928). The source data underlying the statistical results are provided in the Source Data file. Inquiries regarding the dataset can be directed to the corresponding author, Jorie van Haren.

## Code Availability

All analysis scripts and computational modelling code used to process the data and generate the results are publicly available on GitHub (github.com/mesoScopic-Computational-AuditioN-lab). Additional methodological details can be requested from the corresponding author.

## References

[1] Hanne Thielen and Céline R. Gillebert. “Sensory sensitivity: Should we consider attention in addition to prediction?” In: Cognitive Neuroscience 10.3 (2019), pp. 158–160. doi: 10.1080/17588928.2019.1593125.

[2] Clare Press, Peter Kok, and Daniel Yon. “The perceptual prediction paradox.” In: Trends in Cognitive Sciences 24.1 (2020), pp. 13–24. doi: 10.1016/j.tics.2019.11.003.

[3] Rosalyn J. Moran et al. “Free energy, precision and learning: The role of cholinergic neuromodulation.” In: The Journal of Neuroscience 33.19 (2013), pp. 8227–8236. doi: 10.1523/JNEUROSCI.4255-12.2013.

[4] K. Rittershofer et al. “Paradoxical influences of prediction are resolved across time.” In: bioRxiv (2026). Preprint. doi: 10.64898/2026.03.05.709935.

[5] H. H. McDermott et al. “Dissociable dynamic effects of expectation during statistical learning.” In: eLife 13 (2026), RP103689. doi: 10.7554/eLife.103689.

[6] Kechen Zhang and Terrence J. Sejnowski. “Neuronal tuning: To sharpen or broaden?” In: Neural Computation 11.1 (1999), pp. 75–84. doi: 10.1162/089976699300016809.

[7] Arjen Alink, J. Becker, and Helen Blank. “Human vision sequentially prioritizes expected and unexpected information.” In: bioRxiv (Mar. 2026). Preprint. doi: 10.64898/2026.03.23.713176.

[8] Annika Garlichs and Helen Blank. “Prediction error processing and sharpening of expected information across the face-processing hierarchy.” In: Nature Communications 15.1 (2024), p. 3407. doi: 10.1038/s41467-024-47749-9.

[9] Peter Kok, Janneke FM Jehee, and Floris P de Lange. “Less is more: expectation sharpens representations in the primary visual cortex.” In: Neuron 75.2 (2012), pp. 265–270. doi: 10.1016/j.neuron.2012.04.034.

[10] David Richter, Matthias Ekman, and Floris P. de Lange. “Suppressed sensory response to predictable object stimuli throughout the ventral visual stream.” In: The Journal of Neuroscience 38.34 (2018), pp. 7452–7461. doi: 10.1523/JNEUROSCI.3421-17.2018.

[11] Matthew F. Tang et al. “Prediction error and repetition suppression have distinct effects on neural representations of visual information.” In: eLife 7 (2018), e33123. doi: 10.7554/eLife.33123.

[12] Ryszard Auksztulewicz et al. “Rhythmic temporal expectation boosts neural activity by increasing neural gain.” In: The Journal of Neuroscience 39.49 (2019), pp. 9806–9817. doi: 10.1523/JNEUROSCI.0925-19.2019.

[13] David Richter, Micha Heilbron, and Floris P. de Lange. “Dampened sensory representations for expected input across the ventral visual stream.” In: Oxford Open Neuroscience 1 (2022), kvac013. doi: 10.1093/oons/kvac013.

[14] L. Huang et al. “Center-surround inhibition in expectation and its underlying computational and artificial neural network models.” In: eLife 14 (2025), RP107301. doi: 10.7554/eLife.107301.3.

[15] Dirk van Moorselaar and Stefan Van der Stigchel. “Boosting the signal: Expectation-driven gain modulation of preparatory spatial attention.” In: eLife 15 (2026), RP110315. doi: 10.7554/eLife.110315.2.

[16] Benjamin Skerritt-Davis and Mounya Elhilali. “Computational framework for investigating predictive processing in auditory perception.” In: Journal of Neuroscience Methods 360 (2021), p. 109177. doi: 10.1016/j.jneumeth.2021.109177.

[17] Jorie J. G. van Haren et al. “Distinct Roles of Deep and Superficial Cortical Layers in Tone Prediction, Comparison, and Adaptation in Human Auditory Cortices.” In: Nature Communications (2026). doi: 10.1038/s41467-026-75662-w.

[18] Agustín Lage-Castellanos et al. “Investigating the reliability of population receptive field size estimates using fMRI.” In: Frontiers in Neuroscience 14 (2020), p. 825. doi: 10.3389/fnins.2020.00825.

[19] Jae-Jin Kim et al. “An MRI-based parcellation method for the temporal lobe.” In: NeuroImage 11.4 (2000), pp. 271–288. doi: 10.1006/nimg.2000.0543.

[20] Michelle Moerel, Federico De Martino, and Elia Formisano. “An anatomical and functional topography of human auditory cortical areas.” In: Frontiers in Neuroscience 8 (2014), p. 225. doi: 10.3389/fnins.2014.00225.

[21] Eric-Jan Wagenmakers et al. “Assessing model mimicry using the parametric bootstrap.” In: Journal of Mathematical Psychology 48.1 (2004), pp. 28–50. doi: 10.1016/j.jmp.2003.11.004.

[22] Arjen Alink, Hunar Abdulrahman, and Richard N. Henson. “Forward models demonstrate that repetition suppression is best modelled by local neural scaling.” In: Nature Communications 9.1 (2018), p. 3854. doi: 10.1038/s41467-018-05957-0.

[23] Gary Aston-Jones and Jonathan D. Cohen. “An integrative theory of locus coeruleus-norepinephrine function: adaptive gain and optimal performance.” In: Annual Review of Neuroscience 28.1 (2005), pp. 403–450. doi: 10.1146/annurev.neuro.28.061604.135709.

[24] David Richter, Tim C. Kietzmann, and Floris P. de Lange. “High-level visual prediction errors in early visual cortex.” In: PLOS Biology 22.11 (2024), e3002829. doi: 10.1371/journal.pbio.3002829.

[25] Jorie J. G. van Haren et al. “Tonal surprisal and contextual shifts evoke distinct pupil dilation during dynamic sound sequences.” In: European Journal of Neuroscience 63.1 (2026), e70380. doi: 10.1111/ejn.70380.

[26] Jill X. O’Reilly et al. “Dissociable effects of surprise and model update in parietal and anterior cingulate cortex.” In: Proceedings of the National Academy of Sciences 110.38 (2013), E3660–E3669. doi: 10.1073/pnas.1305373110.

[27] Jonathan D. Cohen, Samuel M. McClure, and Angela J. Yu. “Should I stay or should I go? How the human brain manages the trade-off between exploitation and exploration.” In: Philosophical Transactions of the Royal Society B: Biological Sciences 362.1481 (2007), pp. 933–942. doi: 10.1098/rstb.2007.2098.

[28] L. E. Wyatt et al. “Exploration versus exploitation decisions in the human brain: A systematic review of functional neuroimaging and neuropsychological studies.” In: Neuropsychologia 192 (2024), p. 108740. doi: 10.1016/j.neuropsychologia.2023.108740.

[29] Federico De Martino et al. “Frequency preference and attention effects across cortical depths in the human primary auditory cortex.” In: Proceedings of the National Academy of Sciences 112.52 (2015), pp. 16036–16041. doi: 10.1073/pnas.1507552112.

[30] Agustín Lage-Castellanos et al. “Selective attention sharpens population receptive fields in human auditory cortex.” In: Cerebral Cortex 33.9 (2023), pp. 5395–5408. doi: 10.1093/cercor/bhac427.

[31] David H. Brainard. “The Psychophysics Toolbox.” In: Spatial Vision 10.4 (1997), pp. 433–436. doi: 10.1163/156856897X00357.

[32] Denis G. Pelli. “The VideoToolbox software for visual psychophysics: Transforming numbers into movies.” In: Spatial Vision 10.4 (1997), pp. 437–442. doi: 10.1163/156856897X00366.

[33] Yôiti Suzuki and Hisashi Takeshima. “Equal-loudness-level contours for pure tones.” In: The Journal of the Acoustical Society of America 116.2 (2004), pp. 918–933. doi: 10.1121/1.1763601.

[34] Miriam D. Waehnert et al. “Anatomically motivated modeling of cortical laminae.” In: NeuroImage 93.Pt 2 (2014), pp. 210–220. doi: 10.1016/j.neuroimage.2013.03.078.

[35] Matthias Fritsche, Samuel G. Solomon, and Floris P. de Lange. “Brief stimuli cast a persistent long-term trace in visual cortex.” In: Journal of Neuroscience 42.10 (2022), pp. 1999–2010. doi: 10.1523/JNEUROSCI.1350-21.2021.

[36] Johnny van Doorn et al. “Bayesian rank-based hypothesis testing for the rank sum test, the signed rank test, and Spearman’s ρ.” In: Journal of Applied Statistics 47.16 (2020), pp. 2984–3006. doi: 10.1080/02664763.2019.1709053.

[37] Jorie J. G. van Haren et al. Distinct Roles of Deep and Superficial Cortical Layers in Tone Prediction, Comparison, and Adaptation in Human Auditory Cortex. 2025. doi: 10.18112/openneuro.ds006928.v1.0.0. url: https://openneuro.org/datasets/ds006928.

[38] Robert Turner. “How much cortex can a vein drain? Downstream dilution of activation-related cerebral blood oxygenation changes.” In: NeuroImage 16.4 (2002), pp. 1062–1067. doi: 10.1006/nimg.2002.1082.

