## Supplementary materials for "Prediction Dynamically Modulates Gain and Width of Population Frequency Tuning in Human Auditory Cortex"

### A Supplement

#### A.1 Evaluating Dynamic Modulation by Sharpening and Gain Modulation Across Cortical Depth

To investigate whether prediction-elicited dynamic tuning modulations - specifically, sharpening driven by precision-weighted predictions and gain driven by surprisal - are differentially expressed across cortical depth, we partitioned the cortical grey matter within each of the five bilateral regions of interest (Heschl's gyrus (HG), Planum Polare (PP), Planum Temporale (PT), and the anterior and posterior superior temporal gyrus (aSTG, pSTG)) into three depth compartments (deep, middle, and superficial) using an equi-volume approach [34]. To prevent interpolation leaking across depth bins when projecting from anatomical (0.4 mm isotropic) to functional (0.8 mm isotropic) resolution, voxels were assigned to a single depth compartment based strictly on majority volume overlap. Statistical significance across depth compartments ($N = 10$) was evaluated using non-parametric bootstrap $t$-tests ($10^5$ iterations, FDR-corrected) alongside Bayesian Wilcoxon signed-rank tests (Cauchy prior scale $r = \sqrt{2}/2 \approx 0.707$, $10^4$ samples; [36]).

As expected, evaluating the raw cross-validated explanatory power ($R^2$) of all—static and dynamic—models revealed a global increase in variance explained toward the cortical surface across all regions (all layer comparisons $p_{\mathrm{FDR}} \leq 8.0\mathrm{e}{-3}$), reflecting well-documented macrovascular draining patterns typically associated with GE-BOLD [37]. To isolate layer-specific computational effects from this macrovascular confound, we specifically evaluated the relative improvement in explanatory power ($\Delta R^2_{\mathrm{rel}}$) provided by the dynamic models over the static baseline (for a full overview of all laminar comparisons, see Supplementary Table S1).

When evaluating these relative improvements, the superficial gradient disappeared entirely. For the global surprisal-driven gain modulation, the additional variance explained did not differ significantly between deep, middle, and superficial layers across the auditory hierarchy. This lack of a significant laminar effect was consistent in the primary auditory cortex (HG: all layer-pair comparisons $p_{\mathrm{FDR}} \geq 3.5\mathrm{e}{-1}$), the auditory belt (PP: all $p_{\mathrm{FDR}} \geq 1.1\mathrm{e}{-1}$; PT: all $p_{\mathrm{FDR}} \geq 2.1\mathrm{e}{-1}$), and associative regions (aSTG: all $p_{\mathrm{FDR}} \geq 1.1\mathrm{e}{-1}$; pSTG: all $p_{\mathrm{FDR}} \geq 5.6\mathrm{e}{-2}$). Bayesian Wilcoxon signed-rank tests revealed mixed evidence across depth pairs, ranging from anecdotal support for the null hypothesis in primary and belt regions (HG: $BF_{01}$ range 1.37–3.09; PT: $BF_{01}$ range 1.37–1.89) to anecdotal and moderate evidence for the alternative hypothesis in associative areas (aSTG: $BF_{01}$ range 0.26–1.96; pSTG: $BF_{01}$ range 0.14–1.93). Ultimately, the data are inconclusive regarding the presence or absence of a laminar effect for surprisal-driven gain.

Similarly, the relative improvements obtained by the local precision-driven sharpening model were distributed uniformly across cortical depth. We found no significant differences in the relative explanatory gain between layers in the primary auditory cortex (HG: all $p_{\mathrm{FDR}} \geq 3.3\mathrm{e}{-1}$), the auditory belt (PP: all $p_{\mathrm{FDR}} \geq 1.9\mathrm{e}{-1}$; PT: all $p_{\mathrm{FDR}} \geq 1.0\mathrm{e}{-1}$), or associative regions (aSTG: all $p_{\mathrm{FDR}} \geq 1.3\mathrm{e}{-1}$; pSTG: all $p_{\mathrm{FDR}} \geq 3.3\mathrm{e}{-1}$). Consistent with the gain model, Bayesian analyses yielded inconclusive results throughout the temporal cortex, with evidence fluctuating between anecdotal support for the null hypothesis and moderate-to-strong support for the alternative depending on the specific layer comparison (HG: $BF_{01}$ range 0.69–3.00; PP: $BF_{01}$ range 0.52–3.14; PT: $BF_{01}$ range 0.09–2.73; aSTG: $BF_{01}$ range 0.29–2.76; pSTG: $BF_{01}$ range 2.09–3.15).

##### Mechanistic Distinction Between Representational Content and Tuning Modulation

Taken together, the present analyses provided no conclusive evidence for laminar differentiation in dynamic pRF tuning modulation. However, we must be cautious in interpreting these null results, as traditional and Bayesian statistics proved inconclusive. Further research is required to determine the presence or absence of these effects. Evaluating the subtle laminar differences elicited by minor deviations of the dynamic pRFs relative to their static foundation effectively isolated the portions of the signal where the dynamic tuning diverges. Capturing these deviations is inherently constrained by the signal-to-noise ratio of ultra-high-field fMRI, a challenge made worse by the rapid presentation rate of the stochastic tone sequences.

Nevertheless, within the exact same dataset, a different analytical approach - directly modelling the unique representational states of predictions and prediction errors - revealed distinct infragranular and supragranular profiles [17]. Specifically, by partitioning the variance of computational states, our previous work established that expectation-related signals accounted for the most unique variance in deep layers, aligning with the maintenance and updating of the content-specific internal model. Conversely, variance jointly explained by expectations and frequency-specific stimulus preference was strongest in superficial layers, reflecting the tuning-specific alignment of predictions with sensory input.

The seemingly depth-invariant profile of the dynamic pRF modulations observed here contrasts with these established laminar biases. This divergence points to a potential mechanistic distinction; while the representational content of predictions and mismatches relies on distinct laminar streams, their resulting tuning modulation may influence the entire cortical column. In this way, predictive sharpening and gain modulation may act as a layer-agnostic filter, biasing how all cortical layers evaluate and gate incoming signals - both from the top down and from the bottom up - the moment they enter the circuitry, priming the neural population to filter for expected features while universally scaling its sensitivity to environmental shifts.

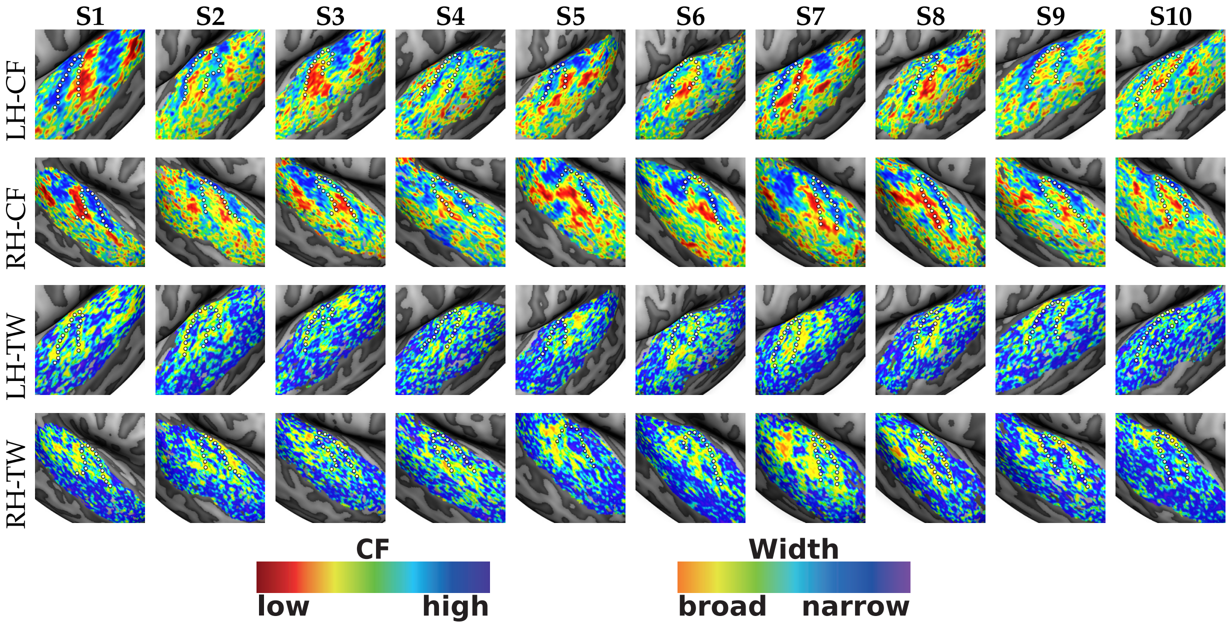


**Figure S1**: **Individual subject pRF centre frequency and tuning width maps.** Rows show estimated population receptive field (pRF) maps for centre frequency (CF) and tuning width (TW), separately for left (LH) and right (RH) hemispheres. Columns correspond to individual subjects (S1–S10). Maps were manually masked to auditory cortex and thresholded at $r > .2$, based on the correlation between voxelwise beta estimates and the fitted Gaussian pRF model. The dotted white line denotes the anatomical border of Heschl's gyrus.

| | Deep vs Middle | | | Middle vs Superficial | | | Deep vs Superficial | | |
|---|---|---|---|---|---|---|---|---|---|
| | $R^2$ | $p_{\text{FDR}}$ | $BF_{01}$ | $R^2$ | $p_{\text{FDR}}$ | $BF_{01}$ | $R^2$ | $p_{\text{FDR}}$ | $BF_{01}$ |
| **Gain (relative)** | | | | | | | | | |
| Heschl's gyrus | 5.6e-5 | 4.3e-1 | 2.93 | -9.9e-5 | 3.5e-1 | 1.37 | -4.3e-5 | 7.4e-1 | 3.09 |
| Planum Polare | 5.8e-5 | 4.3e-1 | 1.98 | -1.6e-4 | 2.4e-1 | 0.33 | -1.0e-4 | 1.1e-1 | 0.83 |
| Planum Temporale | -1.5e-4 | 4.3e-1 | 1.59 | -1.0e-4 | 3.9e-1 | 1.89 | -2.6e-4 | 2.1e-1 | 1.37 |
| anterior STG | -2.2e-4 | 1.2e-1 | 0.26 | -5.2e-5 | 3.9e-1 | 1.96 | -2.7e-4 | 1.1e-1 | 0.52 |
| posterior STG | -8.0e-5 | 4.3e-1 | 1.93 | -2.0e-4 | 5.6e-2 | 0.20 | -2.8e-4 | 6.1e-2 | 0.14 |
| **Sharpening (relative)** | | | | | | | | | |
| Heschl's gyrus | -7.0e-6 | 9.6e-1 | 3.00 | -1.1e-4 | 4.6e-1 | 0.69 | -1.2e-4 | 3.3e-1 | 1.58 |
| Planum Polare | 1.3e-6 | 9.6e-1 | 3.14 | -1.6e-4 | 1.9e-1 | 0.94 | -1.5e-4 | 2.0e-1 | 0.52 |
| Planum Temporale | -1.5e-4 | 1.0e-1 | 0.09 | -5.7e-5 | 6.7e-1 | 2.73 | -2.0e-4 | 3.1e-1 | 0.91 |
| anterior STG | -1.9e-4 | 1.3e-1 | 0.29 | -3.3e-5 | 6.7e-1 | 2.76 | -2.2e-4 | 2.0e-1 | 0.44 |
| posterior STG | 5.4e-6 | 9.6e-1 | 3.15 | -5.4e-5 | 6.7e-1 | 2.09 | -4.8e-5 | 3.3e-1 | 2.29 |
| **Static Baseline (absolute)** | | | | | | | | | |
| Heschl's gyrus | -2.0e-3 | 2.3e-3 | – | -1.9e-3 | 6.5e-4 | – | -4.0e-3 | 4.0e-4 | – |
| Planum Polare | -3.6e-4 | 7.2e-3 | – | -1.2e-3 | 7.2e-3 | – | -1.6e-3 | 4.2e-4 | – |
| Planum Temporale | -1.1e-3 | 7.2e-3 | – | -1.9e-3 | 6.5e-4 | – | -3.1e-3 | 4.0e-4 | – |
| anterior STG | -5.5e-4 | 1.0e-3 | – | -1.3e-3 | 6.5e-4 | – | -1.9e-3 | 4.0e-4 | – |
| posterior STG | -8.5e-4 | 1.0e-3 | – | -8.8e-4 | 6.5e-4 | – | -1.7e-3 | 4.2e-4 | – |
| **Gain (absolute)** | | | | | | | | | |
| Heschl's gyrus | -2.0e-3 | 1.0e-3 | – | -2.0e-3 | 1.4e-3 | – | -4.0e-3 | 5.3e-4 | – |
| Planum Polare | -3.0e-4 | 8.0e-3 | – | -1.4e-3 | 1.8e-3 | – | -1.7e-3 | 8.7e-4 | – |
| Planum Temporale | -1.3e-3 | 8.5e-4 | – | -2.0e-3 | 1.5e-3 | – | -3.3e-3 | 5.3e-4 | – |
| anterior STG | -7.7e-4 | 1.0e-3 | – | -1.4e-3 | 1.5e-3 | – | -2.1e-3 | 5.3e-4 | – |
| posterior STG | -9.3e-4 | 8.5e-4 | – | -1.1e-3 | 2.0e-3 | – | -2.0e-3 | 9.8e-4 | – |
| **Sharpening (absolute)** | | | | | | | | | |
| Heschl's gyrus | -2.0e-3 | 1.7e-3 | – | -2.1e-3 | 1.2e-3 | – | -4.1e-3 | 3.7e-4 | – |
| Planum Polare | -3.6e-4 | 3.8e-3 | – | -1.4e-3 | 5.6e-3 | – | -1.7e-3 | 4.5e-4 | – |
| Planum Temporale | -1.3e-3 | 1.9e-3 | – | -2.0e-3 | 1.7e-3 | – | -3.3e-3 | 3.7e-4 | – |
| anterior STG | -7.4e-4 | 1.9e-3 | – | -1.3e-3 | 1.7e-3 | – | -2.1e-3 | 3.7e-4 | – |
| posterior STG | -8.4e-4 | 3.0e-4 | – | -9.4e-4 | 3.3e-3 | – | -1.8e-3 | 1.5e-3 | – |

**Table S1**: **Absolute and Relative Layer Differences for Static and Dynamic Models.** This table reports depth-resolved statistical results from cross-validated model fits across five auditory ROIs. All values reflect differences in explained variance between cortical depth pairs (deep vs middle, middle vs superficial, deep vs superficial) across participants ($N = 10$). For the relative dynamic models (Gain - driven by global surprisal, and Sharpening - driven by precision-weighted predictions), the $R^2$ difference represents the difference in the relative improvement (dynamic model $R^2$ minus baseline $R^2$) between layers. For the absolute models, it represents the simple difference in raw explained variance between layers. Shown are mean differences with associated paired-sample bootstrapped $p_{\text{FDR}}$ values. Bayesian Wilcoxon Signed-Rank tests ($BF_{01}$) evaluating evidence for the null hypothesis are included for the relative improvement models. $p$-values for depth comparisons are FDR-corrected. Cells with $p_{\text{FDR}} > 0.05$ are shown with a grey background to denote non-significance.

| ROI | $R^2$ | $p_{\text{FDR}}$ |
|---|---|---|
| Heschl's gyrus | 4.2e-3 | <1.0e-4 |
| Planum Polare | 1.1e-3 | 3.0e-3 |
| Planum Temporale | 3.2e-3 | <1.0e-4 |
| anterior STG | 1.4e-3 | 2.0e-4 |
| posterior STG | 1.5e-3 | 2.0e-4 |

**Table S2**: **Cross-Validation of the Static Baseline pRF Model.** This table reports the region-level statistical results for the validation of the static baseline pRF model ($M_0$) evaluated on independent stochastic sequence data from the second scanning session. $R^2$ represents the raw cross-validated variance explained, quantified via leave-one-run-out cross-validation within the second session using static tuning profiles estimated in the first session (controlling for low-level effects of repetition suppression) across five auditory ROIs. Shown are the mean cross-validated $R^2$ values across participants ($N = 10$) alongside their FDR-corrected bootstrapped $p$-values tested against a null baseline of zero.

| ROI | $\Delta R^2_{\text{rel}}$ | $p_{\text{FDR}}$ | $BF_{01}$ | $BF_{10}$ |
|---|---|---|---|---|
| **Gain (Global Surprisal)** | | | | |
| Heschl's gyrus | 0.200 | 3.0e-3 | <0.01 | 87.0 |
| Planum Polare | 0.773 | 3.0e-3 | <0.01 | 201.3 |
| Planum Temporale | 0.319 | 3.0e-4 | <0.01 | 373.4 |
| anterior STG | 0.589 | <1.0e-4 | <0.01 | 351.7 |
| posterior STG | 0.673 | <1.0e-4 | <0.01 | 172.6 |
| **Sharpening (Precision-Weighted Predictions)** | | | | |
| Heschl's gyrus | 0.107 | 2.8e-2 | 0.02 | 46.8 |
| Planum Polare | 0.304 | 1.5e-3 | <0.01 | 289.2 |
| Planum Temporale | 0.134 | 6.0e-4 | <0.01 | 391.4 |
| anterior STG | 0.208 | 1.5e-3 | <0.01 | 349.2 |
| posterior STG | 0.215 | 6.0e-4 | <0.01 | 401.3 |
| **Counterfactual: Gain (Precision-Weighted Predictions)** | | | | |
| Heschl's gyrus | 0.010 | 8.0e-1 | 8.4 | 0.1 |
| Planum Polare | 0.010 | 8.0e-1 | 6.3 | 0.2 |
| Planum Temporale | -0.019 | 9.2e-1 | 9.7 | 0.1 |
| anterior STG | -0.015 | 9.2e-1 | 8.5 | 0.1 |
| posterior STG | -0.050 | 9.2e-1 | 11.9 | 0.1 |
| **Counterfactual: Sharpening (Global Surprisal)** | | | | |
| Heschl's gyrus | -0.017 | 1.0e+0 | 6.6 | 0.2 |
| Planum Polare | 0.002 | 1.0e+0 | 2.7 | 0.4 |
| Planum Temporale | -0.050 | 1.0e+0 | 8.0 | 0.1 |
| anterior STG | -0.080 | 1.0e+0 | 9.1 | 0.1 |
| posterior STG | -0.096 | 1.0e+0 | 8.1 | 0.1 |

**Table S3**: **Relative Improvement of Dynamic Modulations over Static Baseline.** This table reports the region-level statistical results for the relative improvement in cross-validated variance explained when adding dynamic predictive modulations to the static pRF baseline. Results are shown for the primary hypotheses (Gain driven by surprisal and Sharpening driven by precision-weighted predictions) and their counterfactual inverses (Gain driven by precision-weighted predictions and Sharpening driven by surprisal) across five auditory ROIs. $\Delta R^2_{\text{rel}}$ denotes the proportional relative improvement over the static baseline. Shown are the FDR-corrected bootstrapped $p$-values alongside corresponding robust Bayes Factors ($BF_{01}$ and $BF_{10}$) from a Bayesian Wilcoxon Signed-Rank Test quantifying evidence for the null and alternative hypotheses, respectively. Cells with $p_{\text{FDR}} > 0.05$ are shown with a grey background to denote non-significance.

| ROI | $\Delta R^2_{\text{rel}}$ | $p_{\text{FDR}}$ | $BF_{01}$ | $BF_{10}$ |
|---|---|---|---|---|
| **Gain Modulation (Surprisal): Global vs Local** | | | | |
| Heschl's gyrus | 0.217 | 1.5e-3 | <0.01 | 217.8 |
| Planum Polare | 0.757 | 1.0e-4 | <0.01 | 126.6 |
| Planum Temporale | 0.360 | 1.5e-3 | <0.01 | 146.8 |
| anterior STG | 0.634 | 3.0e-4 | <0.01 | 116.5 |
| posterior STG | 0.748 | <1.0e-4 | 0.01 | 99.2 |
| **Sharpening (Precision): Global vs Local** | | | | |
| Heschl's gyrus | -0.058 | 3.8e-1 | 1.3 | 0.8 |
| Planum Polare | -0.242 | 5.9e-3 | 0.03 | 39.5 |
| Planum Temporale | -0.120 | 6.2e-2 | 0.1 | 16.5 |
| anterior STG | -0.183 | 3.6e-3 | <0.01 | 152.2 |
| posterior STG | -0.227 | 3.6e-3 | 0.01 | 90.2 |

**Table S4**: **Relative Improvement of Global vs. Local Predictive Modulations.** This table reports the region-level statistical results contrasting the cross-validated variance explained by global versus local implementations of the dynamic predictive models. A positive $\Delta R^2_{\text{rel}}$ indicates a better fit for the global implementation, while a negative value indicates a better fit for the local implementation. The alternative hypotheses diverge between the two mechanisms: surprisal-driven gain is theoretically hypothesised to operate globally, whereas precision-driven sharpening is hypothesised to operate locally. Consequently, evidence for the alternative hypothesis ($BF_{10}$) follows this hypothesised directionality. Shown are the proportional relative differences, FDR-corrected bootstrapped $p$-values, and corresponding robust Bayes Factors ($BF_{01}$ and $BF_{10}$). Cells with $p_{\text{FDR}} > 0.05$ are shown with a grey background to denote non-significance.
